# DNA-based data storage via combinatorial assembly

**DOI:** 10.1101/2021.04.20.440194

**Authors:** Nathaniel Roquet, Swapnil P Bhatia, Sarah A Flickinger, Sean Mihm, Michael W Norsworthy, Devin Leake, Hyunjun Park

## Abstract

Persistent data storage is the basis of all modern information systems. The long-term value and volume of data are growing at an accelerating rate and pushing extant storage systems to their limits. DNA offers exciting potential as a storage medium, but no practical scheme has been proposed to date that can scale beyond narrow-band write rates. Here, we demonstrate a combinatorial DNA data encoding scheme capable of megabits per second write speeds. The system relies on rapid, combinatorial assembly of multiple smaller DNA parts that are dispensed through inkjet printing. To demonstrate this approach, we wrote approximately 25 kB of information into DNA using our system and read the information back out with commercially available nanopore sequencing. Moreover, we demonstrate the ability to replicate and selectively access the information while it is in DNA, opening up the possibility of more sophisticated DNA computation.

## Introduction

In Nature, DNA is the medium selected for the storage of genetic information. DNA is a promising medium for archival data because of its superior storage density and longevity.^1,2^ It can also be selectively accessed, queried like a database, and even operated on like a parallel computer.^3–5^ Finally, whereas conventional archival media are consistently at the risk of becoming unreadable in the future due to both planned and unintended obsolescence, DNA is guaranteed to remain readable. It will, in fact, become increasingly easier and cheaper to read as DNA sequencing technology improves over time.

While significant advancements have been made in DNA sequencing, the process of writing information into DNA has seen only marginal improvements, due to the fundamental focus on DNA synthesis. Previously published research papers on DNA-based data storage each focus on a particular method of DNA synthesis for the writing of information into DNA and strive to improve the respective synthesis method in order to improve the writing process. Most commonly, other groups have used phosphoramidite chemistry for base-by-base synthesis.^6–8^ Recently, some research groups have explored enzymatic methods for base-by-base synthesis with Terminal deoxynucleotidyl tranferase (Tdt). Though the chemistry of base coupling is different in phosphoramidite synthesis versus Tdt synthesis, both methods use similar multi-step, rate-limiting processes for single nucleotide addition with cycle times around one minute.^9–11^

Our method departs from this previous thinking by considering the most efficient process for the creation of a diverse number of molecules with unique sequences, as well as the encoding scheme that assigns the created molecules to data. Briefly, we create the entropy by assembling pre-fabricated DNA fragments rather than by traditional base-by-base synthesis. This approach circumvents current bottlenecks, shifting a cyclical chemistry problem to a mechanical challenge, solvable with current inkjet printing technologies. In the system we have developed, one module rapidly deposits DNA, while another module activates the chemistry for assembly, and a final module combines all assembled DNA into a pooled library. All together, the instrument is capable of megabits per second write speeds. We have also developed a novel encoding scheme to use these molecules for the storage of information.

## Results

### Symbol Generation and Encoding Methodology

A set of DNA fragments (Components) are designed and partitioned into a fixed number of disjoint sets (Layers). These Components are prefabricated using conventional DNA manufacturing techniques such as base-by-base synthesis. Longer DNA molecules (Identifiers) are assembled from these Components by ligating them together, where only one Component from each Layer is chosen per Identifier, and the Layers have a pre-defined fixed order. Because the Components are designed with overhangs such that they will only ligate to members of adjacent Layers, the Layer that a Component belongs to determines the relative position of each Component in the Identifier. We call this the Cartesian Product of Components (CPC) method of entropy generation. FIG 1A schematically illustrates an example of a Component library. Each Component comprises a unique central region and two edge sequences that are common to all Components within a Layer. The central regions are designed to not only to be able to robustly identify each Component despite error-prone sequencing, but also functionally for hybridization such that each Component forms stable duplex along its central region, and additionally such that each Component contains a unique primer binding site for targeted selection of Identifiers, which is used for random access. The distal edge sequences on all Components of the first and last Layers, termed terminal sequences, are designed to comprise primer binding sites which are used for various steps downstream of sequencing, including clean-up, sequencing preparation, and replication. Besides the terminal sequences, all other edge sequences are single-strand overhangs designed to facilitate specific and efficient ligation between Components of adjacent Layers.

**FIG 1.**
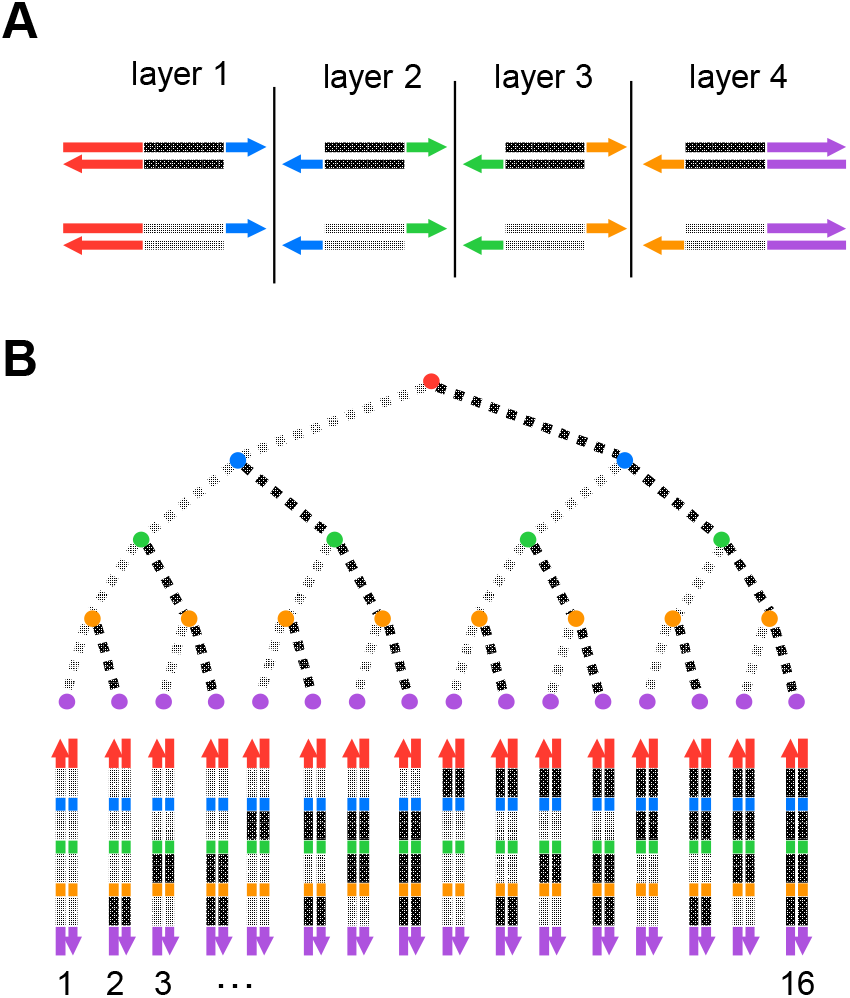
Schematic of a Component library. A. DNA Components have unique central regions and two edge sequences that are common to a layer. B. An Identifier is an assembly of Components, one DNA selected from each layer. In this example, 16 unique Identifiers are assembled using 8 Components.

We define an “Identifier space” as the set of all possible Identifiers that can be constructed from a particular library of starting Components. Generally, the size of an Identifier space grows exponentially with the number of Layers and is given by the formula 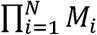 where *N* is the number of Layers and *M*_*i*_ is the number of Components in the *i*-th Layer. Each Component can be specified by a coordinate (*i, j*) where *i* is its Layer and *j* is the Component number ranging from *1* to *M*_*i*_. FIG. 1B illustrates an example of an Identifier space of size 16 made from a Component library of eight Components partitioned into four Layers, with two Components per Layer (represented in FIG 1A). We organize the Identifiers in an ordered tree data structure, which we call a C-tree. Each level of the C-tree corresponds to a Layer *i*, with all nodes of the level having *M*_*i*_ branches arranged in a fixed order, each corresponding to a Component of the Layer. Extending the branch ordering, the Identifiers in the C-tree can also be ordered from 0, 1, …, *N* − 1 by enumerating each root-to-leaf path in the C-tree with a post-order traversal of the tree, respecting the branch order at each level. The logical position of any given Identifier, which we call the rank of the Identifier, is then defined by counting the number of Identifiers that precede the Identifier. We use the rank of each Identifier to encode the address of a bit of information, either a *1* or a *0*. To encode a given string of bits *b*_0_, *b*_1_, … *b*_*N*-1_, we assemble and pool the Identifiers that encode the positions of bits with a value of *1*. That is, we only assemble and pool the identifier of rank *i* if and only if *b*_*i*_ = 1. This scheme may be extended to encode with alphabets of any non-binary arity.

### Writer System Overview

A Component set of 110 Components partitioned into 15 Layers of five Components each and one Layer of 35 Components provides a C-tree with an Identifier space sufficient to encode a terabit-length bit string. The total number of ligation reactions and the total number of liquid transfers required to collocate the reactions is approximately 30 × 10^9^ reactions and 9 × 10^11^ transfers, respectively. Assuming 1000 transfers are completed per second by off-the-shelf liquid handling robots, a terabit collocation would take approximately 30 years. To alleviate this intractability, we designed a writer capable of high throughput collocation using an ink-jet printing approach. In our system, Identifiers are synthesized in a three-step process: (1) collocation of Components into individual reaction spots, (2) incubation to allow ligation, and (3) pooling to collect the resultant Identifiers. We built a specialized DNA writing system with separate hardware modules for each of these steps: a print engine, an incubator, and a pooler (FIG 2A). The modules are physically connected by a chassis that threads a continuous polypropylene sheet (webbing) through each of the modules providing a surface on which to create assembly reaction spots. The webbing speed is variable, and experiments reported here used a speed of 16 m/min.

**FIG 2.**
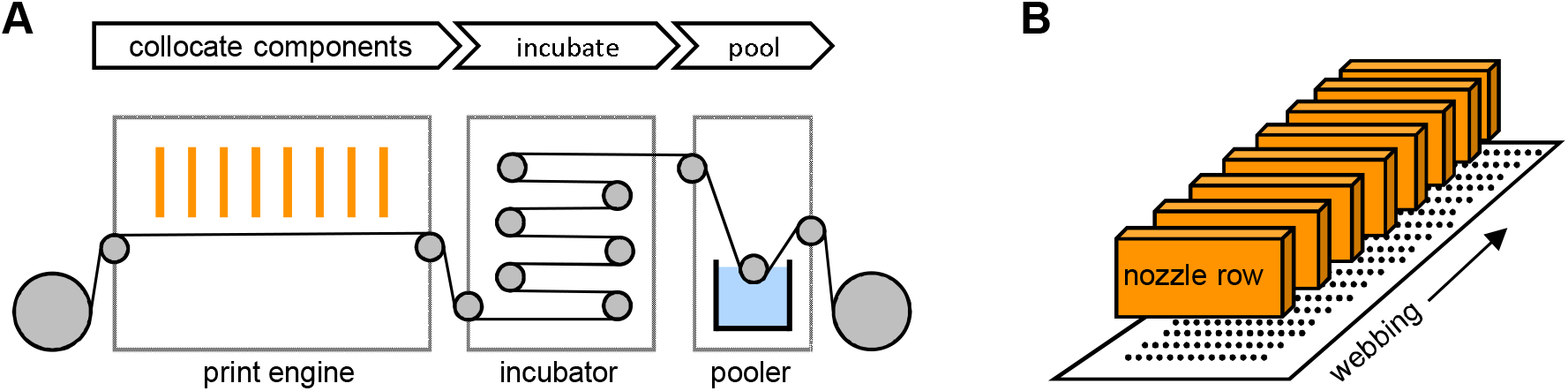
DNA writer schematic. A. The DNA writer has 3 critical functions: collocation of DNA Components using a print engine module, reaction spot incubation using an incubator, and reaction spot pooling using a pooler. B. The print engine module contains an array of inkjet printheads that dispense DNA components.

The print engine contains an array of inkjet printheads with a total of 120 nozzle rows. FIG 2B provides a top-down schematic of a section of the print engine. Each nozzle row is connected to an individual ink supply and can print droplets on demand. We have formulated inks for dispensing Components, as well as ligase and buffer. The inks containing ligase and buffer are also plumbed to 6 nozzle rows, enabling the writer to accommodate starting libraries of up to 114 Components. Each nozzle row is aligned such that any combination of inks can be overprinted onto the webbing as it passes underneath and perpendicular to each nozzle row. Thus, the print engine is designed to generate reaction spots on the webbing, where each reaction spot contains a programmable combination of inks configured to form at least one Identifier.

After the print engine, the webbing and the freshly printed reaction spots enter the incubator module for enzymatic assembly. The webbing travels a spiral path length through a series of rollers before exiting the incubator. In our current roller configuration, a maximum incubation time of 150 seconds is achieved when the web is running continuously at 16 m/min, but longer incubations can be accommodated by intermittently pausing the web inside of the incubator. Upon exiting the incubator, the webbing passes through the pooler module, which includes a sonicated reservoir of an EDTA solution to simultaneously collect reaction spots and inhibit enzymatic ligation. The pooler is designed for continuous collection of reaction spots over the course of hours in liter-scale volumes. For smaller write jobs, the pooler module can be bypassed, and the reaction spots may instead be manually excised, pooled, and washed with mL-scale volumes of 50 millimolar EDTA solution.

### Print Runs

To validate our method and system, we encoded and decoded three compressed text files selected from Wikipedia. The articles for “Meselson-Stahl experiment” (File 1), “Timeline of information theory” (File 2), and “*Encyclopedia Galactica*” (File 3) combined to 8580 bytes. We encoded the files into an Identifier space constructed from a Component library of 16 Layers and 44 total Components, with one Layer having 12 Components, two Layers containing three Components, and every other Layer having two Components each. FIG 3A illustrates a full path of the C-tree used to order the Identifier space. The levels of the C-tree begin with Layer 16, then Layer 1, then Layer 15, then Layer 2, and so on, alternating back and forth while moving inward from the two ends of the Identifier. This logical structure makes it possible to access the data contained in any range of Identifiers with sequential PCR reactions. Starting with a PCR reaction using primers that anneal to Components on Layer 1 and Layer 16, each subsequent iteration of PCR utilizes primers that anneal to specific Components on the outer Layers of the amplicon from the previous iteration, thus enabling top-down tree traversal to any range of bits.

**FIG 3.**
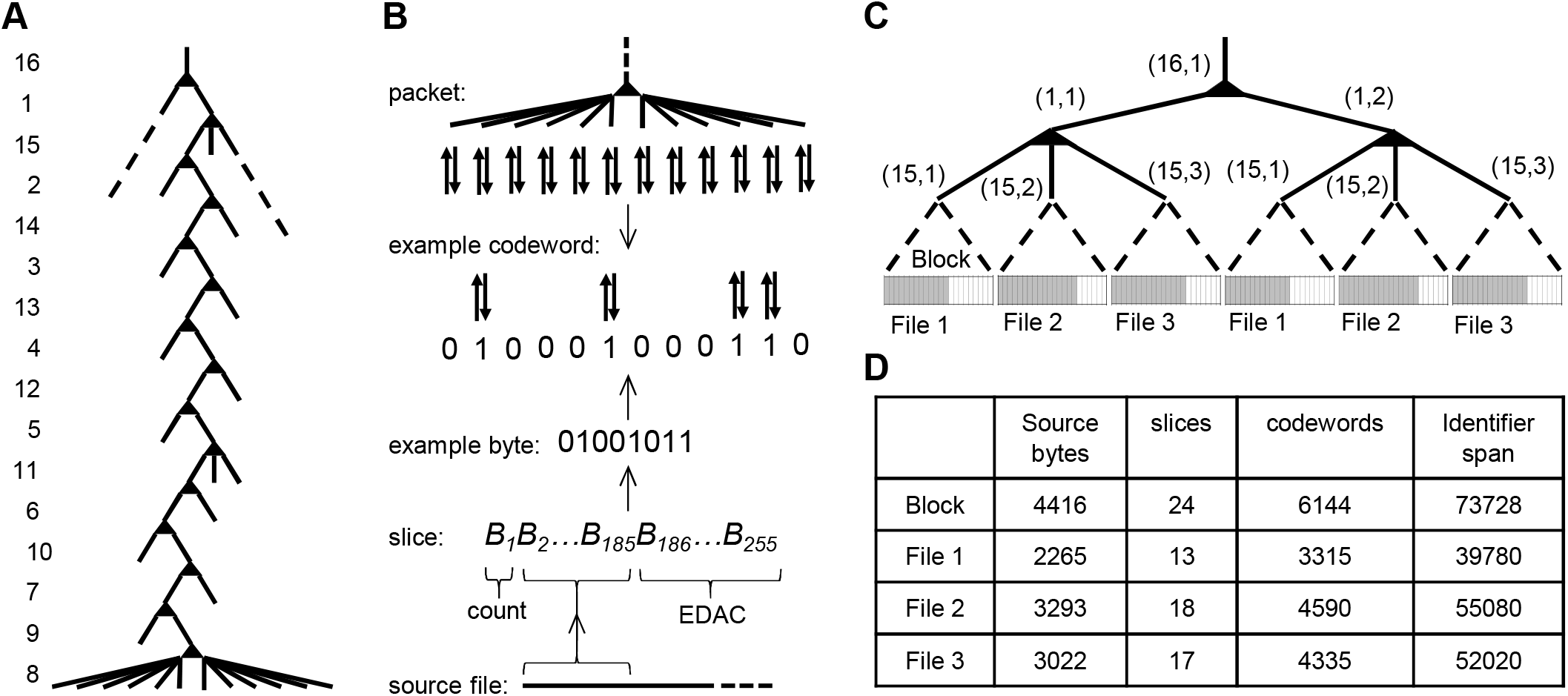
Encoding digital information into DNA Identifiers. A. Illustration of the C-tree used to order Identifier space. B. Demonstration of the logical process for encoding each file. C. Example of packet allocation of the Identifier space. D. Overview of file size in relation to the block capacity.

FIG 3B demonstrates the logical process for encoding each file. As an initial encoding step, each source file was broken up into slices and padded with additional error detection and correction (EDAC) bytes using Reed-Solomon codes. Specifically, we used slices of length 255 bytes with 70 EDAC bytes. Additionally, one byte on each slice was allocated to count the number of source bytes on the slice. All slices had 184 source bytes, except for the last slice of each file which had the remainder of file source bytes up to 184. Next, slice bytes were translated to binary codewords of length 12 and weight 4. The codeword length of 12 was chosen to match the final fan-out of our tree data structure, which makes it so that the Identifiers that make up a codeword always share all Components in common except for the Components of a single Layer, in this case Layer 8. When writing the data, we programmed each reaction spot to multiplex the assembly of all Identifiers that encode a codeword, in this case 4, by dispensing one Component from each of the Layers leading up to Layer 8 on the C-tree, and then dispensing 4 Components from Layer 8. The fixed weight constraint enabled uniform volumes and concentrations across all reactions. The value of 4 was chosen because it is the smallest *k* such that 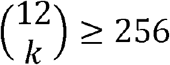, the minimum entropy needed to encode a byte.

The full size of our Identifier space was enough to store 73,728 bytes, and our files, after being encoded into slices, aggregated to 12,240 bytes. Thus, we encoded each file twice for a total encoded data size of 24,480 bytes. FIG 3C shows how the files were allocated to packets of the Identifier space. The packets were defined by Identifiers that share a common prefix on the C-tree to make them easily accessible with PCR. Though in general, packets need not have a fixed size, here we used packets defined by the first three levels of our C-tree, resulting in packets capable of holding up to 24 slices each – more than enough space to accommodate each individual file. FIG 3D provides an overview of the size of each file in relation to the capacity of the blocks.

The data was written using 46 nozzle rows, one for each of the 44 Components, and one nozzle row for ligase ink and one nozzle row for buffer. Every reaction spot was programmed to encode a codeword and accordingly was created on the web by overprinting combinations of 19 droplets comprising Components (4 droplets from Layer 8 and one droplet from each of the other 15 Layers) and one droplet each of ligase and buffer, for a total of 21 overprinted droplets. Each droplet was approximately 7 pL, giving a total reaction spot volume of 147 pL. Quality control images were taken of all of the reaction spots as they traveled from the print engine to the incubator (FIG 4A). The reaction spots were printed with a pitch of approximately 330 µm. Reaction spot radii were observed to form a tight normal distribution with a mean of 52 µm and standard deviation of 0.083 µm, indicating precise print volume deposition and overprinting. Every codeword was printed six times, yielding a total of 146,880 reaction spots for the assembly of 97,920 unique Identifiers printed at an approximate rate of 125,000 reaction spots per second. This translates to an encoding rate of approximately 1 megabit per second, as each reaction encodes a byte (8 bits).

**FIG 4.**
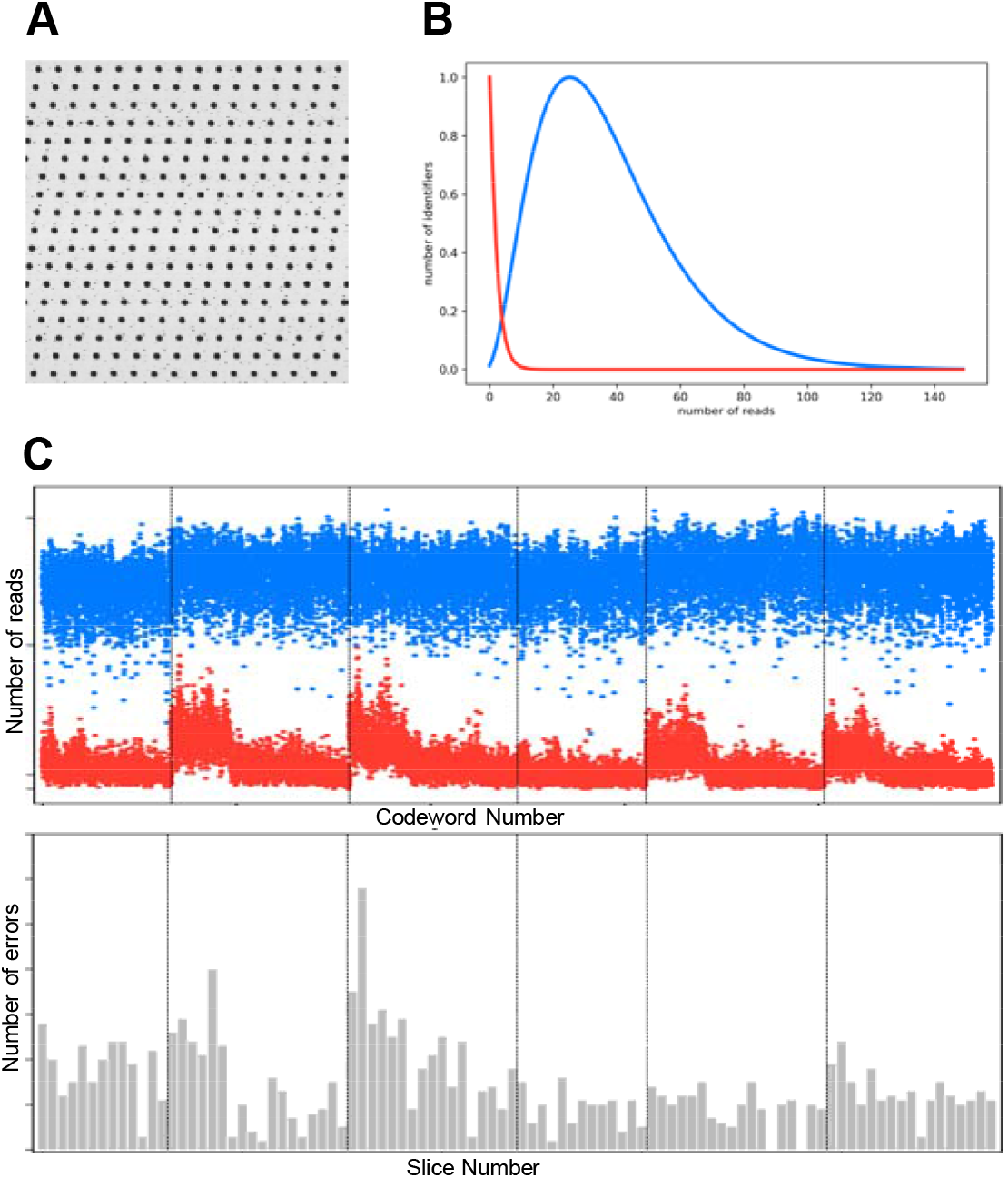
Summary of reaction spots produced by print engine. A. Image of reaction spots as part of in-process quality control. B. Distribution of true Identifiers (blue) and false Identifiers (red) detected by sequencing. C. The top panel shows true Identifiers (blue) compared with false Identifiers (red) per codeword. The bottom panel compares the number of errors per slice. The top of the panel represents the maximum number of correctable errors.

The reaction spots were incubated for 10 minutes and then excised from the webbing and pooled together in DNA Cleanup Binding Buffer (New England Biolabs). A 528 base pair full-length assembly product (an Identifier) was separated from shorter assembly products using gel extraction (Thermo Fisher Scientific E-Gel). The pooled sample with isolated full-length Identifiers are classified as the “root sample”. We subsequently decoded the information by (1) sequencing the root sample on a nanopore sequencing device (Oxford Nanopore MinION), (2) aligning the sequences to our Identifier library, and (3) using the recovered Identifiers to translate the data back to the original slice bytes. In total, we sequenced approximately 4 million reads corresponding to Identifiers and observed at least one copy of all but 26 of the 97,920 Identifiers that we intended to make, termed “true Identifiers”.

We identified two error modes that may result from our DNA writer. Missing true Identifiers in a codeword could lead to a “byte erasure” in which a byte is not decodable because not all four Identifiers comprising the codeword were observed by sequencing. Alternatively, we may also observe copies of some Identifiers that we did not intend to make, termed “false Identifiers”. The presence of false Identifiers in a codeword could lead to a “byte substitution” in which a byte value is misinterpreted due to the observation of one or more incorrect Identifiers within the codeword. When more than 4 Identifiers are observed for a codeword, we select the 4 Identifiers observed at the highest number of times in sequencing. If no such determination can be made, then a byte erasure results. If a false Identifier is observed more times than a true Identifier, then a byte substitution results. When all true Identifiers are observed at a higher count than any of the false Identifiers, then the correct byte is decoded.

FIG 4B. shows that the distribution of true Identifiers and false Identifiers in the sequencing data are well resolved. Although false Identifiers were detected in the sequencing data, their counts were sufficiently low to be easily distinguishable from their true counterparts. We define signal-to-noise ratio, or SNR, as a metric to quantify the separation of the true and false Identifier distributions and calculate it as the mean observed count of true Identifiers divided by the mean observed count of false Identifiers. The overall observed SNR was 22.1. The top panel of Fig 4C illustrates a trace of the mean true Identifier copies versus the mean false Identifier copies per codeword. Generally, the true Identifiers maintain steady levels and good separation from false Identifiers. The bottom panel of FIG 4C illustrates a trace of the number of errors per slice, with the top of the trace window corresponding to the error tolerance of our Reed-Solomon code. Using Reed-Solomon, we calculated errors as the sum of the number of erasure errors and 2x the number of substitution errors. Although there is a fairly uniform incidence of errors throughout the dataset and an average error rate of 0.133 per byte, all the errors are correctable because they fall far below the tolerance of our error protection encoding (for this experiment we used 28% Reed-Solomon error correction bytes).^12^

### Replication and Random Access

Along with the capability to encode information into DNA, we demonstrated the compatibility of our writing with performing replication and selective access. For replication, we amplified our root sample using PCR with primers complementary to the common terminal sequences of our Identifiers. An aliquot of the root sample estimated to contain 1000s of copies per true Identifier underwent 8 thermal cycles for an approximate 256-fold amplification. The replicated sample was replicated again 3 additional times in series to test the robustness of the encoded information against successive rounds of the replication protocol. Similarly, for random access, we performed PCR on the root sample using primers that anneal to Components (1,1) and (16,2) to access the Identifiers contained in the latter half of our Identifier space (See Fig. 3A). As with the root sample, we used nanopore sequencing to decode replication and random access samples. Table 1 summarizes the quality of the decoded information from each sample. The error rate for all samples were sufficiently low for decodability (See Table 1).

**Table 1.** Quality of decoded information.

| Sample | Percent decoded | Reads per byte | Erasure rate | Mutation rate | Corrected error rate | SNR |
| --- | --- | --- | --- | --- | --- | --- |
| root | 100 | 162 | 0.021 | 0.056 | 1.8E-17 | 22.1 |
| psg1 | 100 | 165 | 0.015 | 0.012 | 9.8E-23 | 44.6 |
| psg4 | 100 | 82 | 0.041 | 0.024 | 2.0E-11 | 43.9 |
| ram | 100 | 123 | 0.017 | 0.011 | 1.0E-23 | 55.1 |

## Discussion

We demonstrated the ability to write information into DNA at megabit per second write speeds using our system. Though the demonstration wrote a small aggregate amount of information, around 25 kB (2 copies of each of 3 files) in a single run, the system is configured to write up to 140 GB in one day at its maximum speed and capacity. The system is able to achieve this write speed by using a novel writing method, CPC, that circumvents the traditional chemistry bottleneck of base-by-base synthesis in the writing process. In our system, the major determinant of write speed is the speed at which assembly reactions are created. To this end, we leveraged technology from the inkjet printing industry to build our system. Standard inkjet printheads are capable of dispensing inks at kilohertz rates from hundreds of nozzles in parallel, enabling millions of reactions to be set up per second. Top-of-the-line inkjet printheads on the market currently have thousands of nozzles per nozzle row and fire at rates greater than 10 kHz, which may facilitate near gigabit per second write speeds, a critical milestone towards commercial viability.^13,14^ Further, inkjet printhead technology is still improving and will continue serving as a positive externality for writing systems that invoke our method. Regardless, the CPC method itself can be readily deployed on other liquid handling platforms.

Our CPC method coupled with our encoding scheme fundamentally differs from other DNA-based storage work reliant on base-by-base synthesis for writing data in DNA. Specifically, using prefabricated DNA Components for writing data in DNA dramatically reduces DNA cost because large stores of Components can be made and used over multiple print runs and eliminates synthesis time required for base-by-base synthesis. Additionally, the encoding method presented herein encodes data in the specific subset of Identifiers chosen, rather than in the sequence of the molecules themselves. This creates new opportunities for the design of error correcting codes and data structures. Unlike the base-by-base synthesis approaches, these code-level designs need not be constrained by sequence-level considerations such as the length of our molecules or the avoidance of certain sequences. We have demonstrated that the writing method is compatible with replication and random access, both critical building blocks for future explorations of DNA-based computation. Further, because information is not written at single base-resolution, our system has a larger tolerance for sequencing errors. This opens an avenue for the development of faster, cheaper sequencing instruments that would otherwise be unsuitable for genomic applications but are tenable for our method of data storage. Thus, our system not only demonstrates a dramatic increase in writing speed for DNA data storage but also has potential for higher read speeds, enabling both commercially-viable archival DNA data storage and future highly-parallelized computations in DNA.

## Methods

### Annealing

Custom DNA oligonucleotides with 5’ phosphate modifications were ordered from Thermo Fisher Scientific, Inc. (Waltham, MA, USA). Complementary oligonucleotide pairs were mixed at an equimolar ratio with 50 mM KCl and annealed at 90°C for 10 minutes, followed by incubation at 4°C for a minimum of 12 hours before use. Annealing efficiency was confirmed by visualizing the dsDNA on a 4% EX Agarose E-Gel (Thermo Fisher Scientific Inc., Catalog Number G401004; Waltham, MA, USA).

### Pooling DNA identifiers

Once writing was completed, the web motion was stopped and the relevant portion of the web was excised with scissors. The relevant sector of the webbing was trimmed with a clean scalpel to roughly 5 cm wide and inserted into a 50 mL polypropylene tube with 5 mL of New England Biolabs Monarch DNA Cleanup Binding Buffer (Ipswich, MA, USA) amended with 10 mM EDTA. The tube was rotated several times to wash the buffer over the reaction drops and collect the identifiers in the buffer. Pooled identifiers were purified with New England Biolabs Monarch DNA Cleanup Kit columns (T1030; Ipswich, MA, USA) per the manufacturer’s protocol and eluted in 20 µL nuclease-free water.

### Size Selection

Shorter assembly products were removed through a gel extraction of the 528-bp full length assembly product. The purified pooled product was loaded onto 2% EX Agarose E-Gel (Thermo Fisher Scientific Inc.; catalog number G401004; Waltham, MA, USA) at a mass of <200 ng per lane and ran on the Invitrogen E-Gel Power Snap Electrophoresis System for 10 minutes (Waltham, MA, USA). Excised bands with purified with New England Biolabs Monarch DNA Gel Extraction Kit (T1020; Ipswich, MA, USA) per the manufacturer’s protocol and eluted in 20 µL nuclease-free water. The mass of the size-selected, purified product was quantified with qPCR. qPCR reactions were 10 µL and contained 1x KAPA Sybr Fast Low-Rox qPCR Master Mix (KK4621; Roche Sequencing, Pleasanton, CA, USA) with 200 nM each primer. qPCR reactions were run on a Q5 QuantStudio (ThermoFisher, Waltham, MA, USA). The cycling protocol consisted of a 3 minute denaturing step at 95°C, followed by 40 cycles of a 2-step PCR stage (3s at 95°C and 1 min at 60°C) and a melt curve stage consisting of 1s at 95°C, 20s at 60°C and 1s at 95°C. A quantitation standard consisting of a 528bp double stranded gBlock Gene Fragment ordered from Integrated DNA Technologies, Inc. (Coralville, Iowa, USA) with analogous primer binding sites to the identifiers was run in triplicate at 5 concentrations (386 fg/µL, 38.6 fg/µL, 3.86 fg/µL, 0.386 fg/µL, 0.0386 fg/µL). Triplicate no-template controls contained 1x KAPA Sybr Fast Low-Rox qPCR Master Mix (KK4621; Roche Sequencing, Pleasanton, CA, USA) with 200 nM each primer and water only. The size-selected, purified product was amplified to a concentration between 1-10 ng/µL in order to generate sufficient mass for sequencing. 50-µL PCR reactions contained 1x Phusion HF buffer (New England Biolabs; Ipswich, MA, USA), 200 µM dNTPs, 200 µM forward primer, 200 µM reverse primer, 1 unit Phusion DNA Polymerase (New England Biolabs; Ipswich, MA, USA), and 30-50 pg template. Amplification was performed on a 3×32 well Proflex PCR system (Applied Biosystems; Waltham, MA, USA). Reactions were amplified with a multi-step thermocycler protocol, consisting of a 30 second denaturing step at 98°C, followed by 19 cycles of 10s at 98°C, 10s at 60°C and 20s at 72°C, followed by 5 minute final extension at 72°C. PCR reactions were purified with Monarch DNA Cleanup Kit columns (T1030; New England Biolabs; Ipswich, MA, USA) per the manufacturer’s protocol and eluted in 20 µL nuclease-free water.

### Sequencing

DNA amplicons were quantified on a Qubit 4 Fluorometer (Invitrogen; Waltham, MA, USA) using the Qubit 1x dsDNA HS assay kit (Q33231; Waltham, MA, USA). Libraries were prepared with the Oxford Nanopore Adaptors by Ligation library prep kit (SQK-LSK109; Oxford, England). Library quantitation was performed on a Qubit 4 Fluorometer (Invitrogen; Waltham, MA, USA) using the Qubit 1x dsDNA HS assay kit (Q33231; Waltham, MA, USA) prior to loading on the flowcell. Sequencing was performed on the Oxford Nanopore Minion device with a SpotOn flowcell (R9.4.1; Oxford, England).

## Acknowledgements

The authors would like to thank Tracy Kambara, David Turek, and Anne Condon for their critical review of the manuscript. The authors would also like to thank Milena Lazova for her early work on the system.

## Notes

### Competing Interest Statement

Authors are employees or former employees of Catalog Technologies.

